# Comparative analysis of dengue and Zika outbreaks reveals differences by setting and virus

**DOI:** 10.1101/043265

**Authors:** Sebastian Funk, Adam J. Kucharski, Anton Camacho, Rosalind M. Eggo, Laith Yakob, Lawrence M. Murray, W. John Edmunds

## Abstract

The pacific islands of Micronesia have experienced several outbreaks of mosquito-borne diseases over the past decade. In outbreaks on small islands, the susceptible population is usually well defined, and there is no co-circulation of pathogens. Because of this, analysing such outbreaks can be useful for understanding the transmission dynamics of the pathogens involved, and particularly so for yet understudied pathogens such as Zika virus. Here, we compared three outbreaks of dengue and Zika virus in two different island settings in Micronesia, the Yap Main Islands and Fais, using a mathematical model of transmission dynamics, making full use of commonalities in disease and setting between the outbreaks. We found that the estimated reproduction numbers for Zika and dengue were similar when considered in the same setting, but that, conversely, reproduction number for the same disease can vary considerably by setting. On the Yap Main Islands, we estimated a reproduction number of 8.0–16 (95% Credible Interval (CI)) for the dengue outbreak and 4.8–14 (95% CI) for the Zika outbreak, whereas for the dengue outbreak on Fais our estimate was 28–102 (95% CI). We further found that the proportion of cases of Zika reported was smaller (95% CI 1.4%–1.9%) than that of dengue (95% CI 47%–61%). We confirmed these results in extensive sensitivity analysis. They suggest that models for dengue transmission can be useful for estimating the predicted dynamics of Zika transmission, but care must be taken when extrapolating findings from one setting to another.

**Author Summary:** Dengue and Zika are related viruses and are both transmitted by mosquitoes. Dengue is well described and affects people around the world. Zika, on the other hand has only recently caused outbreak in human populations, and it has been suggested that its behaviour might similar dengue. To investigate this, we compared three outbreaks of dengue and Zika in island populations of the pacific: two dengue outbreaks and one Zika outbreak. Island outbreaks are useful laboratories for understanding the spread of infections because they are usually short, well-identified episodes, whereas elsewhere it can be difficult to identify the properties of outbreaks when different viruses spread at the same time. In our investigation of the outbreaks in Micronesia we found that dengue and Zika virus did, indeed, behave similar in outbreaks they caused on the Yap Main Islands. A dengue outbreak on the smaller island of Fais, on the other hand, was different from the dengue outbreak on Yap in that transmission seems to have been more efficient. We conclude that dengue outbreaks are indeed a good model for Zika outbreaks when considered in the same setting, but that one must be careful when comparing outbreaks in different settings.

## Introduction

Many infections of humans are transmitted by mosquitoes. Dengue virus is one of the major pathogens infecting humans worldwide, causing an estimated 50–100 million cases resulting in about 10,000 deaths annually [1]. Confined mainly to tropical regions because of its reliance on transmission through *Aedes* mosquitoes, it is endemic in more than 150 countries across the world [2]. Its four circulating serotypes cause a wide range of clinical symptoms and severities, with most cases resolving without progressing to the more severe forms, dengue hemorrhagic fever or dengue shock syndrome. Upon infection following bite by an infectious female mosquito, the virus undergoes a period of incubation before progressing to disease in an estimated 20-50% of infected people [3,4], with symptoms lasting approximately one week. The relative infectiousness of symptomatically and asymptomatically infected people remains a topic of active study, with recent evidence indicating that symptom-free people are more infectious to mosquitoes than clinically symptomatic people [5,6]. Infection results in lifelong immunity to same serotype but subsequent infection with heterologous serotypes is associated with higher rates of severe dengue [7].

Zika virus, a member of the *Flaviviridae* family like dengue and also transmitted by *Aedes* mosquitoes was discovered in Africa in 1947 [8]. Formerly believed to be mostly confined to primate species, it has caused occasional cases in humans across Africa and equatorial Asia in the decades after its discovery, before causing its first observed outbreak in humans on the Yap Main Islands, Micronesia, in 2007 [9,10]. Following further outbreaks on Pacific islands in 2013/14 [11–13], cases of an illness characterised by skin rash were reported from Brazil beginning in March 2015 and Zika virus circulation confirmed in May 2015 [8, 14,15]. Zika virus appears to largely cause asymptomatic infection or mild disease and a non-itchy rash. However, it has recently been linked to neurological issues in rare cases, particularly microcephaly when contracted in pregnancy [16] and Guillain-Barré syndrome [17, 18]. A recent increase in reported occurrences of microcephaly in Brazil has led to the declaration of a Public Health Emergency of International Concern by the World Health Organization, to “reduce infection with Zika virus, particularly among pregnant women and women of childbearing age.” [19].

In contrast to dengue, Zika virus has not been described in great detail, and its epidemiology in human populations remains poorly understood. Here, we characterise the epidemiology of dengue and Zika outbreaks in tropical island settings by comparing three outbreaks in Yap State, Micronesia: the outbreak of Zika virus on the Yap Main Islands in 2007, a dengue outbreak on the Yap Main Islands in 2011, and a dengue outbreak on the island of Fais. Island outbreaks are a particularly useful vehicle for understanding transmission dynamics as cases usually occur in episodic outbreaks, limiting interaction between pathogens and reducing the chances of misclassification. Moreover, all three outbreaks share particular characteristics: the two dengue outbreaks share the infecting agent; the two outbreaks on the Yap Main Islands the setting; and the Zika outbreak on the Yap Main Islands and the dengue outbreak on Fais that they probably struck immunologically naϊve populations. Moreover, evidence suggest that both *Aedes aegypti* and *Aedes hensili* are important epidemic vectors in both settings, with the latter only recently having been implicated in outbreaks of arboviruses [20, 21]. We exploit these relationships to comparatively study the three outbreaks by simultaneously fitting a mathematical model to all three time series, holding the parameters constant between the outbreaks where they represent a common element.

## Methods

### Outbreak setting

Yap State is one of the four states of the Federal States of Micronesia, consisting of the Yap Main Islands (also called Yap Proper or simply Yap) and fourteen outer atolls spanning an area of approximately 120 km^2^. The Yap Main Islands consist of four major inhabited islands and six smaller ones that form a contiguous land mass of approximately 79 km^2^. The 7,370 inhabitants of the Yap Main Islands (2010 census, population density 93/km^2^) live in villages, the largest of which is the capital of Yap State, Colonia (population 3,126), with the remaining villages mostly located along the shore line. Fais is one of the outer islands of Yap State which lies about 270 km to the East of the Yap Main Islands and has a much smaller land mass (2.6 km^2^). The population of 294 (2010 census, density 113/km^2^) is concentrated in a single population centre that spans approximately a quarter of the island’s area (Fig. 1).

**Figure 1.**
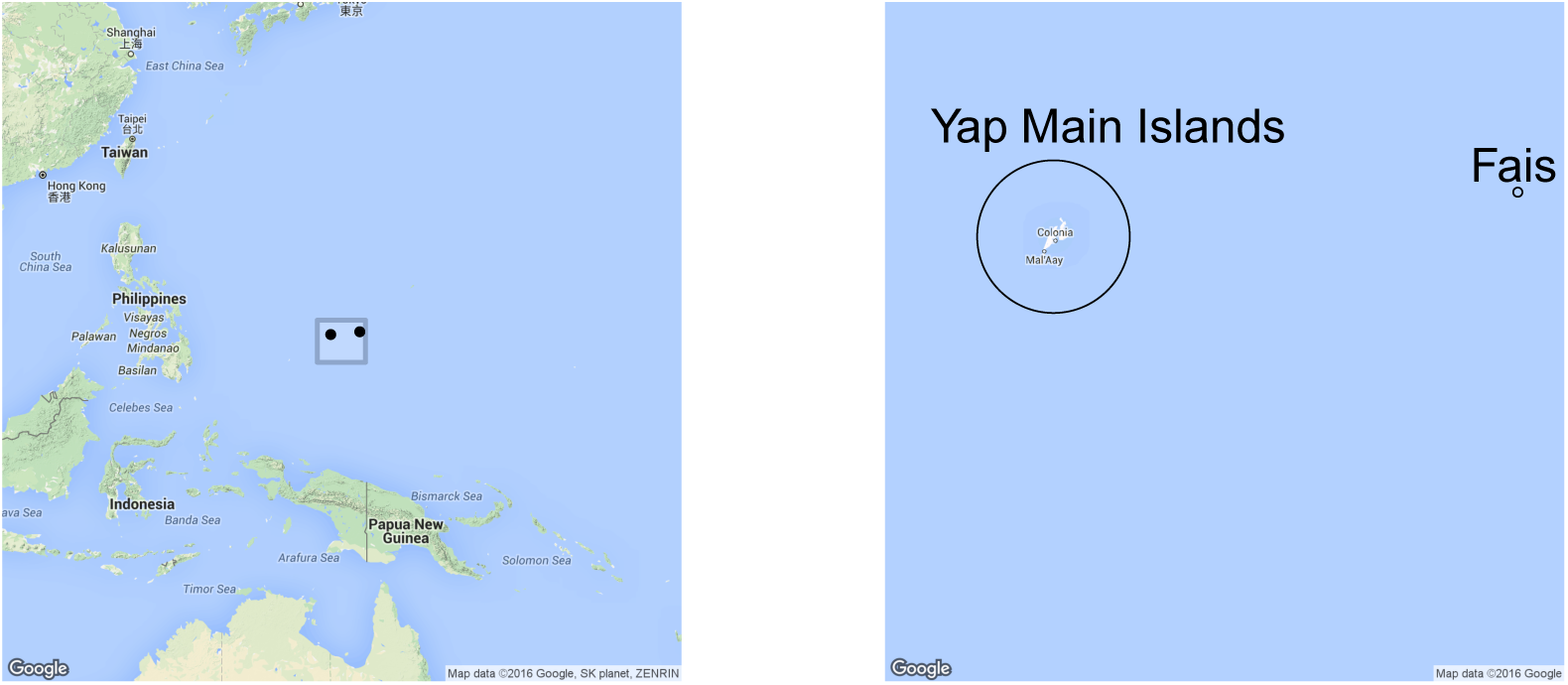
Geographical location of the Yap Main Islands and Fais. The two islands are marked in the left panel with black dots, and shown in more detail on the enlarged map in the right panel.

The Yap Main Islands have experienced several outbreaks of dengue in the past, including an outbreak of serotype 4 in 1995 [20] and an outbreak of serotype 1 in 2004 [22]. The outbreak of Zika in 2007, on the other hand, was the first observed outbreak of Zika overall [9]. The outbreak of dengue in Fais, too, is believed to have been the first ever on the island [23].

Because of its stable climate, mosquito numbers are not believed to vary seasonally in Micronesia [24].

### Data

The dengue time series from the Yap Main Islands and Fais consist of suspect dengue cases as identified by the Yap Department of Health [23]. Clinically suspected dengue cases were identified using the WHO (2009) case definition. A small proportion of cases (9%) were reported on outer islands and included in the time series for the Yap Main Islands as we did not have access to a time series where the two were separated. Dengue virus serotype 2 was confirmed by reverse transcriptase polymerase chain reaction by the CDC Dengue Branch, Puerto Rico. The Zika time series from the Yap main islands consists of probable and confirmed cases as identified in a combination of prospective and retrospective surveillance at all health centres on Yap [9].

All three time series of cases are shown in Figure 3 and summarised in Table 1. The outbreak of Zika on the Yap Main Islands had its first cases reported with onset in mid-April 2007 and the last in July 2007. Overall, a total of 108 cases were classified as probable (59) and confirmed (49) in a population of 7,370, and 73% (95% CI 68–77) were later found with evidence of recent Zika infection in a household survey [9]. The outbreak of dengue on the Yap Main (and Outer) Islands began with a case with disease onset on 1 September, 2011, and two more onsets on the following day. The next case was reported with onset a week later, on 8 September, followed by another cluster around 15 September, and sustained spread beginning another week later, around 22 September, 2011. The peak of the outbreak occurred in the week beginning 24 November, 2011, with 142 cases reported with onset during that week. The last cases were reported with onset on 16 February, 2012.

**Figure 3.**
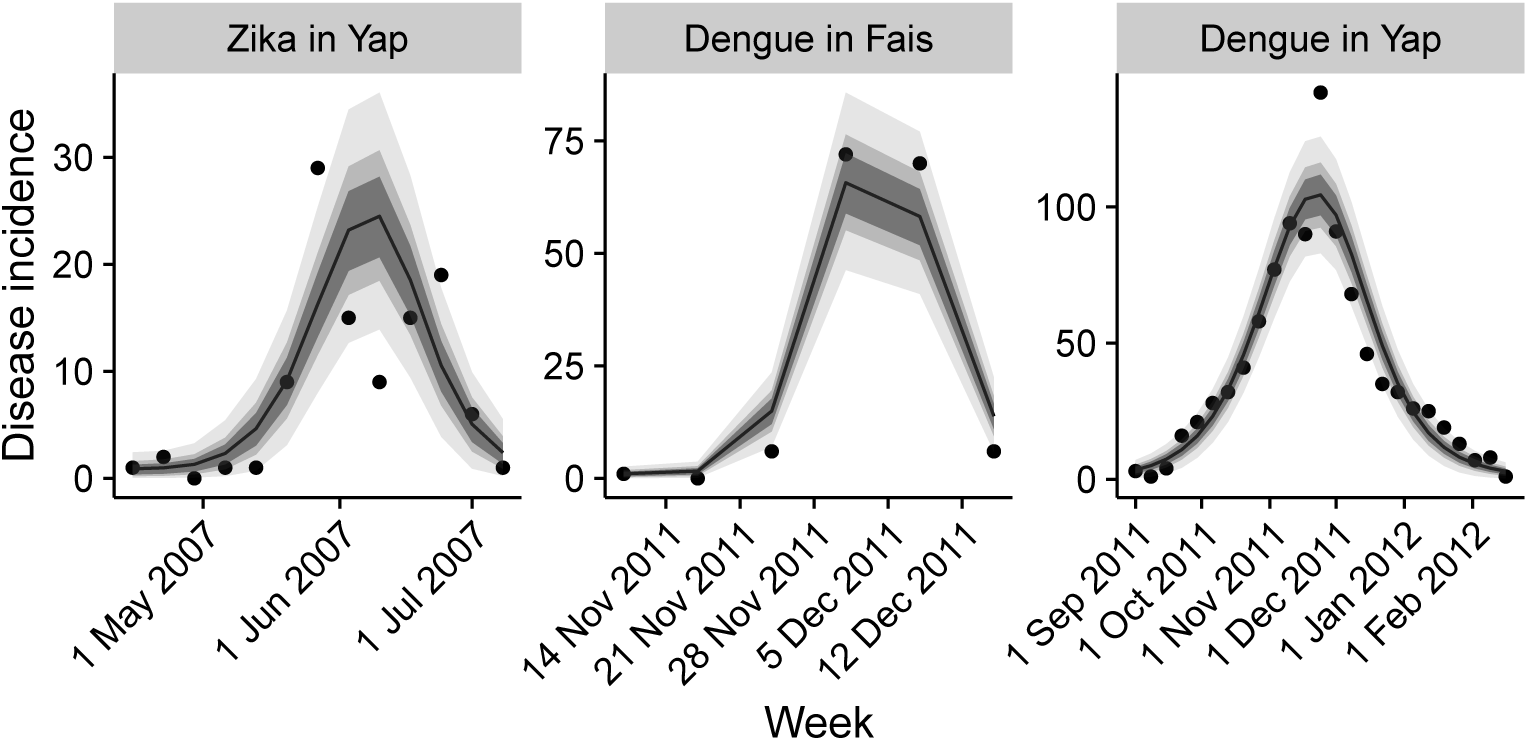
Time-lines of the outbreaks. Left to right: Zika virus on the Yap Main Islands, 2007; dengue outbreak on the Yap Main (and Outer) Islands, 2011 and Fais, 2011. Shown are the data (weekly incidence) as dots, and model fits (median, line; interquartile range, dark grey; 72% and 95% credible intervals, lighter shades of grey).

**Table 1.** Outbreak characteristics. Summaries of the three outbreaks.The outbreak of dengue on Fais overlapped with the outbreak on the Yap Main Islands. It began on 10 November, 2011, with onset of disease in the likely index case. No further case was reported for 16 days, before cases started increasing after the second reported case (onset on 27 November, 2011) to a peak of 72 cases reported with disease onset in the week beginning 1 December, 2011. The last reported disease onsets were 2 cases on 20 December, 2011. Overall, 157 clinical cases were reported among the 294 residents.

| Location | Disease | Population | Reported cases | Duration (weeks) |
| --- | --- | --- | --- | --- |
| Yap | Zika | 7370 | 108 | 13 |
| Yap | Dengue | 7370 | 978 | 24 |
| Fais | Dengue | 294 | 155 | 6 |

### Transmission model

We implemented a variant of the Ross-McDonald model [25,26], schematically depicted in Fig. 2. The human population of size N_H_ was divided into susceptible (S_H_), incubating or exposed (E_H_), infectious (I_H_) and recovered (R_H_) compartments. The mosquito population of unknown size was divided into the proportion susceptible (s_M_), incubating (e_M_) or and infectious (i_M_). We assumed that the size of the human (N_H_) and vector populations did not vary over the course of the modelled outbreaks (i.e., we ignored birth and death rates in the human populations and assumed them to be the same in the vector populations), and that the symptomatic period in humans agrees with the infectious period [27]. We further assumed that infection results in immunity that lasts for at least the duration of the outbreak, and that vertical transmission in the mosquito population can be neglected [28].

In our model, everybody who gets infected can transmit the virus to mosquitoes [6]. Any lack of symptomatic disease is reflected in the mean proportion of cases reported *r*, as defined in the likelihood function below. The system of ordinary differential equations (ODEs) governing the outbreaks are:

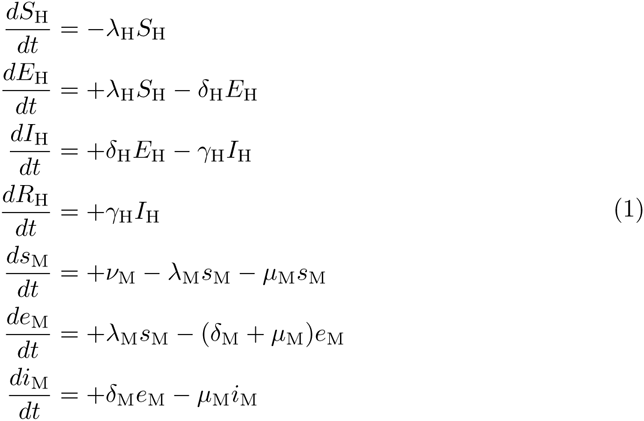

**Figure 2.**
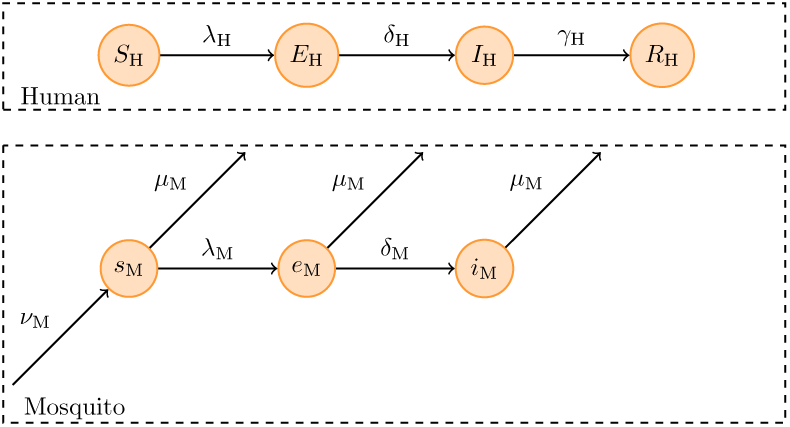
Model structure. Only compartments that are relevant to the observed case series are depicted. For details of the parameter values, see text.

Here, λH and λM are the forces of infection acting on humans and mosquitoes, respectively, δ_H_ = 1/D_inc,H_ and *δ*_*M*_ = 1/*D*_inc,M_ are the incubation rates, defined as the inverse of the average incubation periods *D*_inc,H_ and *D*_inc,M_ in humans and mosquitoes, respectively, γH = 1/*D*_inf,H_ is the recovery rate in humans, defined as the inverse of the average duration of infectiousness, *ν*_M_ is the birth rate of female mosquitoes or number of susceptible female mosquitoes born per female mosquito per unit time, here assumed to be equal to the mosquito death rate µ*M* = 1/*D*_life,M_, defined as the inverse of the average mosquito life span *D*_life,M_. This ensured that mosquito population sizes remained constant over the course of each outbreak.

The forces of infection can be written as

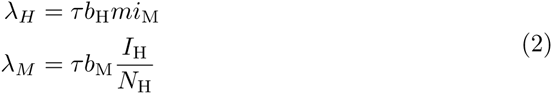

where τ is the number of human blood meals taken by a single female mosquito per unit time, b_H_ and b_M_ are the probabilities that a bite by an infectious female mosquito leads to infection in a human and a bite on an infectious human leads to infection in a mosquito, respectively, and m is the number of female mosquitoes per human.

The human-to-human reproduction number of this model is

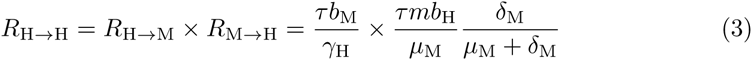

The basic reproduction number of the system, or the average number of secondary infections (in human or mosquito) from a primary infectious bite can be calculated from the next-generation matrix [29], and is the square root of the human-to-human reproduction number given in Eq. 3.

### Generation Intervals

The equilibrium generation interval, or the mean time between the infection of a primary case and its secondary cases, relates reproduction numbers (which only describe reproduction per generation, without an explicit time scale) to the time scale of transmission. For our model, in an equilibrium situation it would be [30]:

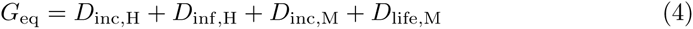

In an outbreak situation, observed generation intervals deviate from the theoretical value at equilibrium and change over time. When new infections are generated at approximately exponential rate, observed mean generations interval are smaller than the equilibrium value as most infectious people will only just have been infected [31]. This issue has recently been generalised to the whole distribution of generation intervals, and beyond assumptions of exponential growth [32].

For Zika, the generation interval has been estimated to be between 10 and 23 days [33], combining estimates for *D*_inc,H_ of 3–12 days, *D*_inc,M_ of 4–6 days, assuming *D*_inf,H_ = *D*_life,M_ = 0, that is that mosquitoes are infected by humans and vice versa just after their infectious period started, as well as an additional delay before symptomatic humans become viraemic of 3–5 days. If humans are, instead, taken to be viraemic for the first 3–5 days from symptoms onset [34], the estimated range shortens to 7–18 days. This should be taken as a lower limit for observed generation intervals, as in reality some infections will be caused by humans/mosquitoes that have been infectious for some time.

A second study that estimated the equilibrium generation interval using all the components of Eq. 4 and drawing from a systematic review of the natural history of the infection [35]. Assuming that humans or mosquito are equally likely to cause infection in mosquitoes or humans, respectively, the generation interval was estimated to be 20 days (mean, 95% CI 15.6–25.6), with a standard deviation of 7.4 days (mean, 95% CI 5.0 – 11.2), using an average mosquito life time of 5 days with standard deviation of 1.7 days [36].

### Parameter estimation

To fit the model to the data sets, we used a Bayesian framework, generating samples from the posterior distribution using Markov-chain Monte Carlo (MCMC). The observation likelihood at each data point was assumed to be distributed approximately according to a Poisson distribution with rate *rZ*_H_, where *Z*_H_ is the number of new human infections of Zika per reporting interval *r* is the proportion of these infections that get reported, estimated using a normal approximation with mean and variance both equal to *rZ*_H_. We only had access to a weekly time series of Zika on the Yap Main Islands, and therefore aggregated the daily time series of dengue cases to weekly numbers to make estimates comparable between time series.

We fixed the biting rate to 1 per day [37]. Since we did not have enough information on mosquito life span to inform a full prior distribution, we further fixed the life span of the mosquito to either 1 week [36] or 2 weeks [38], and compared the two sets of fits using the Deviance Information Criterion or DIC [39]. We modelled the other natural history parameters (intrinsic and extrinsic incubation periods and infectious period in humans) with dengue-like priors, assuming that infectiousness starts 1.5 days before symptom onset [36, 40] and ends 1.5 days before their end. These prior distributions overlap with ones that have recently been estimated from the available data for Zika virus infections [35, 36].

We estimated the remaining parameters of the model by fitting to all three time series simultaneously, with the following constraints: probabilities of infection from a potentially infectious bite, proportion reported, intrinsic and extrinsic incubation periods and human infectious periods were all to be disease-specific but the same across settings; mosquito densities, on the other hand were to be setting-specific but the same across the two pathogens, reflecting potential differences in the sizes of vector populations but also in human population density and behaviour.

For the outbreak of dengue the Yap Main Islands, we assumed that only a proportion *q* of the population was susceptible to infection. For the Zika outbreak on the Yap Main Islands, we assumed that the whole population was susceptible to infection. In other words, our Zika model is the assumed equivalent of a single-serotype dengue model not incorporating cross-reactivity between heterologous viruses or serotypes. The dengue outbreak in Fais, too, was assumed to strike a fully susceptible population, as it was the first known outbreak of dengue on the island. All outbreaks were started with a single infectious case, and date at which that case became infectious fitted as a separate parameter (rounded to the week) for all three outbreaks.

The MCMC procedure for parameter estimation was implemented using the *libbi* software package [42], run from the statistical package *R* [43] using the *RBi* [44] and *RBi.helpers* [45] packages. After adapting the size and shape of the multivariate normal proposal distribution in trial runs, the algorithm was run for 10 million iterations and convergence confirmed visually. All code and data used to generate the results are available at http://github.com/sbfnk/vbd.

### Alternative models

We fitted two modified models to a data set containing an additional data point included in the fit to reflect the final outbreak size observed in a serological study on the Yap Main Islands [9]. The likelihood at this data point was given by a normal distribution centred around the final size, with a standard deviation of 2.2% to reflect the 95% confidence interval reported in the serological study. In one model, the population size of Yap Main Islands would be reduced by a factor *ρ* [41], whereas in the other one the initial proportion susceptible would be a proportion *q* of the whole population but everybody susceptible to mosquito bites, as in our model for the dengue outbreak on the Yap Main Islands.

We further fitted a two-patch metapopulation model to the outbreaks on the Yap Main Islands. While we did not have any spatially resolved data to inform such a model, the outbreak of Zika on the Yap Main Islands could be interpreted to consist of two peaks, a structure that would be expected to reproducible by a two-patch model. In this model, the outbreak starts in a patch which contains a proportion ϕ of the total population. This and another patch share the same parameters, and human in each patch exert a force of infection on mosquitoes in the other (representing human movement) that is reduced by a factor *σ* with respect to the force of infection within each patch.

## Results

The models with mosquito life spans of 1 week vs 2 weeks fit the data equally well (DIC difference < 1). Assuming that both were equally likely to be true and combining the posterior distributions, the estimated disease-specific durations of infection and incubation largely corresponded to the given prior distributions (Table 2). There was, however, a more than twenty-fold difference in the proportion of infectious people reported, between a median estimate of 53% (IQR 51–56, 95% CI 47–61) for dengue and 1.6% (IQR 1.5–1.7, 95% CI 1.4–1.9) for Zika. Location-specific parameters indicated a considerable difference in the number of female mosquitoes per person, with a mean estimate of 1.0 (IQR 0.69–1.5, 95% CI 0.38–8.4) on the Yap Main Islands and 4.7 (IQR 3.4–7.2, 95% CI 2.1–30) on Fais. The proportion of the population initially susceptible to dengue on the Yap Main Islands was estimated to be 27% (IQR 26–29, 95% CI 24–32).

**Table 2.** Posterior means, 95% credible intervals (CIs) and prior distributions of estimated parameters. Yap: Yap Main Islands. Parameters given for the distributions are the lower and upper bound for (Log-)uniform distributions, and mean and standard deviation for (Log-)normal distributions. Durations are given in units of days and rates in unite of days^−1^. CI: credible interval.

| Disease-specific parameters |  |  |  |  |  |  |
| --- | --- | --- | --- | --- | --- | --- |
|  | Dengue |  | Zika |  | Prior | Reference |
|  | Median | 95% CI | Median | 95% CI |  |  |
| $D_{\text{inf,H}}$ | 4.2 | (3.7, 4.5) | 4.4 | (4.0, 4.8) | Normal(4.5, 1.75) | [46] |
| $D_{\text{inc,H}}$ | 4.7 | (1.6, 7.6) | 4.9 | (2.2, 7.7) | Normal(4.4, 0.25) | [27, 40] |
| $D_{\text{inc,M}}$ | 8.6 | (4.3, 15) | 9.1 | (4.4, 17) | Normal(6.5, 1.15) | [27] |
| $b_{\text{H}}$ | 0.63 | (0.17, 0.98) | 0.59 | (0.14, 0.97) | Uniform(0,1) | n/a |
| $b_{\text{M}}$ | 0.77 | (0.16, 0.99) | 0.58 | (0.07, 0.98) | Uniform(0,1) | n/a |
| $r$ | 0.53 | (0.47, 0.61) | 0.016 | (0.014, 0.019) | Uniform(0,1) | n/a |
| Location-specific parameter |  |  |  |  |  |  |
|  | Yap |  | Fais |  | Prior | Reference |
|  | Median | 95% CI | Median | 95% CI |  |  |
| $m$ | 1.0 | (0.38, 8.4) | 4.7 | (2.1, 30) | Log-uniform(0.1, 100) | n/a |
| $\tau$ | 1 | – | 1 | – | fixed | [37] |
| $q$ | 0.27 | (0.24, 0.32) | | | Uniform(0, 1) | n/a |
| Common parameter |  |  |  |  |  |  |
|  | Mean | 95% CI |  |  | Prior | Reference |
| $D_{\text{life,M}}$ | 7 or 14 | | | | fixed | [36, 38] |

The median estimates of the human-to-human reproduction number, R_H→H_ were 11 (IQR 9.7–13, 95% CI 8.0–16) for dengue on the Yap Main Islands, 7.6 (IQR 6.3–9.6, 95% CI 4.8–14) for Zika on the Yap Main Islands, and 51 (IQR 40 – 71, 95% CI 28–102) for dengue on Fais (Fig. 4). By combining the estimated parameters between settings and disease, we estimated R_0_ for Zika on Fais to be 35 (posterior mean, IQR 26–52, 95% CI 18–79). The differences in *R*_0_ between Yap and Fais are reflected in the different estimated differences in the number of female mosquitoes per person, which results in differences in the number of bites experienced per person.

**Figure 4.**
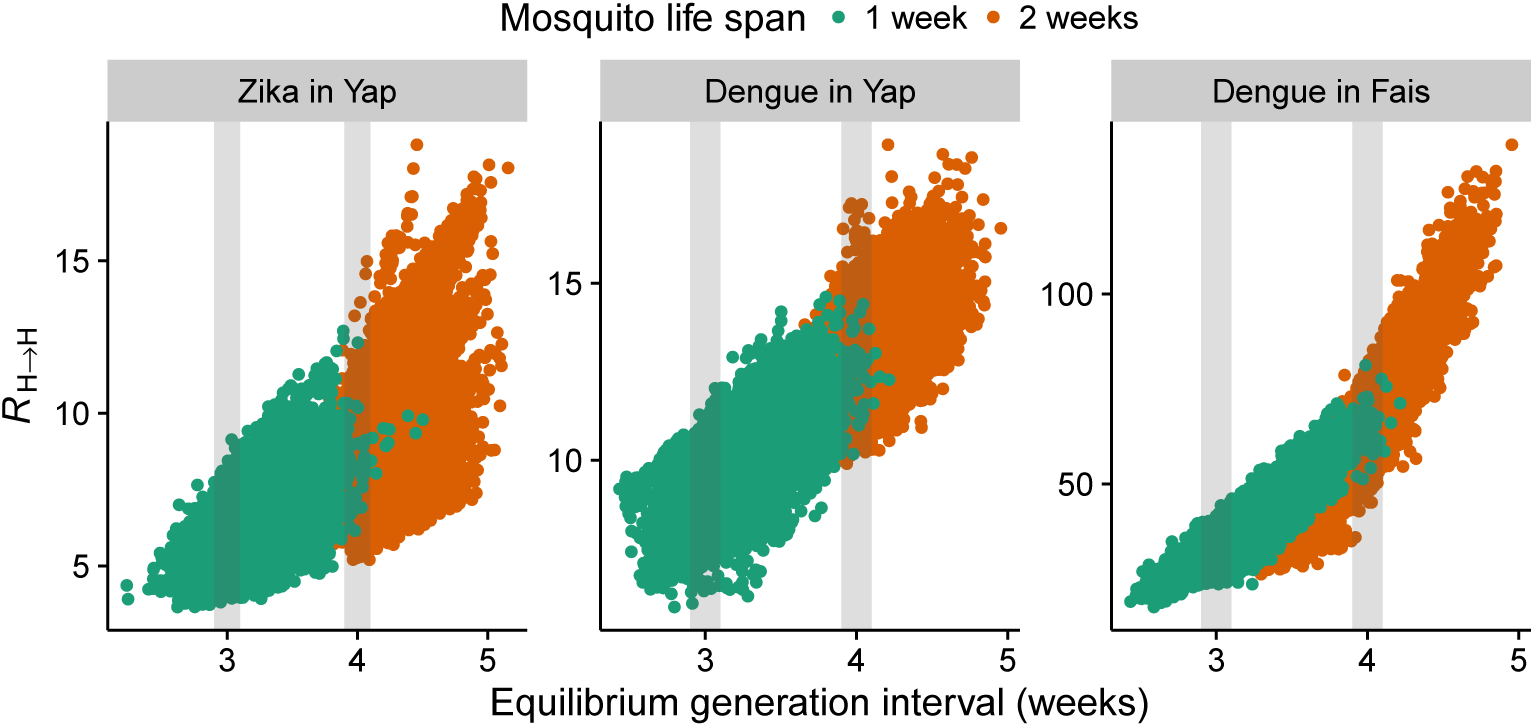
Relationship between the human-to-human reproduction number and the equilibrium generation interval. Human-to-human reproduction number *R*_H→H_ and equilibrium generation interval *G*_eq_ in posterior samples split by whether mosquito life spans *D*_life,M_ was 1 week (green) or 2 weeks (brown). Regions used to estimate the reproduction number in Table 3 are shaded in grey.

Much of the variation in *R*_0_ is explained by the different lengths of the generation interval which was poorly identified from the data (Fig. 4, Table 3). This is particularly the case for dengue in Fais, where all infections occurred in one or two generations, depending on the length of the generation interval (Fig. 5).

**Table 3.** Posterior mean, IQR and 95% credible interval (CIs) of the human-to-human reproduction number for different generation intervals(± 0.1 weeks) from samples of the posterior distribution.

| Disease | Setting | Generation interval (weeks) | $R_{\text{H} \rightarrow \text{H}}$ | | |
| --- | --- | --- | --- | --- | --- |
|  |  |  | median | IQR | 95% CI |
| Zika | Yap | 3 | 5.7 | (5.2,6.3) | (4.4,7.6) |
|  |  | 4 | 8.4 | (7.5,9.5) | (6.2,12) |
| Dengue | Yap | 3 | 9.0 | (8.4,9.6) | (7.4,11) |
|  |  | 4 | 13 | (12,14) | (11,15) |
| Dengue | Fais | 3 | 34 | (31,36) | (27,41) |
|  |  | 4 | 65 | (60,70) | (51,79) |

**Figure 5.**
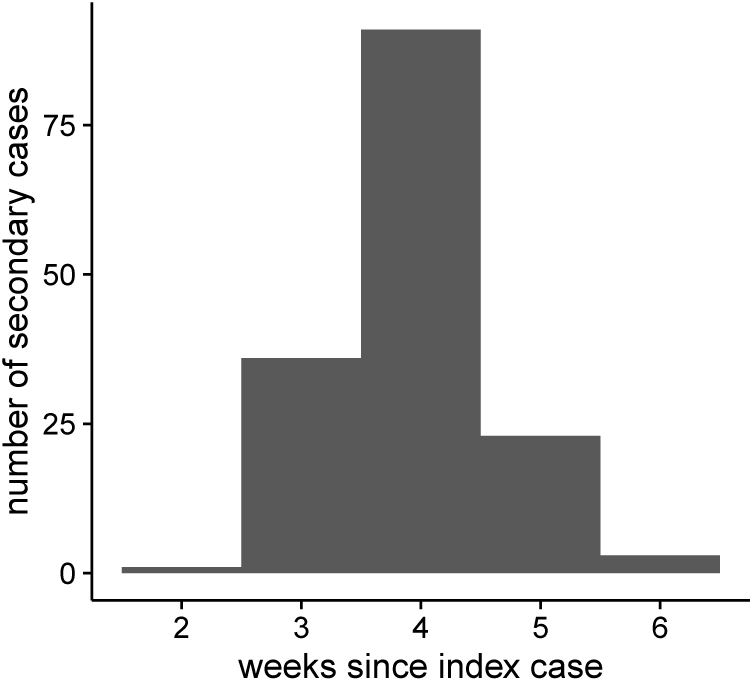
Distribution of secondary cases for the dengue outbreak in Fais. Time is measured in weeks since the index case (all symptom onset).

The alternative models with reduced population size or reduced susceptibility against Zika on the Yap Main Islands were both able to reproduce the observed proportion infected of 73% (see Supporting Information S1 Text). In the model with reduced population size the initial proportion susceptible to dengue on the Yap Main Islands was estimated to 37% (median, IQR 35–39, 95% CI 32–44), leading to a smaller human-to-human reproduction number of 8.7 (median, IQR 7.3–10, 95% CI 6.0–13) and greater proportion of Zika cases reported of 2.2% (median, IQR 2.1–2.3, 95% CI 1.9–2.7). In the model where only a proportion of the population q was susceptible to infection with Zika on the Yap Main Islands, the estimate of the proportion susceptible to dengue and human-to-human reproduction numbers were unchanged, while the proportion of Zika cases reported increased to 2.2% (median, IQR 2.1–2.4, 95% CI 1.8–2.7) The two models described the data equally well (DIC difference < 1). The alternative two-patch metapopulation model produced very similar parameter fits to the single-patch model. In particular, the fit to the outbreak of Zika on the Yap Main Islands produced a single peak unless it was viewed in isolation.

## Discussion

We have analysed three outbreaks of mosquito-borne disease on small islands of Micronesia using a mathematical model. We exploited the overlap between those outbreaks in setting and disease to constrain parameter values and used this to investigate differences in transmission dynamics. While we found large difference between the reproduction numbers for dengue in two different island settings, our estimates of the reproduction numbers for dengue within the same settings are similar.

Our approach of fitting three time series concurrently and with common parameters helped identify some parameters that would not be identifiable by observing the outbreak in isolation. The proportion of cases of dengue reported was informed by the final size of the dengue outbreak in Fais which, in turn, enabled estimation of the initial proportion susceptible of the dengue outbreak on the Yap Main Islands, again from the final outbreak size. With these two parameters established, the reproduction number of dengue in the two settings could be estimated from initial growth rate and outbreak duration, as a function of the generation interval. The generation intervals themselves were poorly identified in the data, and the corresponding marginal posterior distributions largely overlapping with the prior distributions.

Parameters for the Zika outbreak on the Yap Main Islands were similarly identified. With the reproduction number given by the initial growth rate and outbreak duration, the proportion of cases reported could be estimated from the reported final outbreak size of the epidemic. In this context it should be noted that with the values of the reproduction number we estimated, one would expect nearly all of the population to get infected, in contrast to the 73% (68–77) estimated to have been infected in a serological study after the outbreak [9]. It remains an open question how to best reconcile a rapidly growing epidemic that spreads through large parts of a population in a few generations without rendering everybody seropositive, a phenomenon also observed in the 2013–14 Zika outbreak in French Polynesia [13,41,47]. In the case of Zika on the Yap Main Islands, there might be several reasons for the discrepancy between modelled outbreak sizes and observed serology, such as the sensitivity of the used diagnostic test or lack of seroconversion at low-level exposure. If, on the other hand, the measured seropositivity reflects true infection history, its discrepancy with our modelled outbreak sizes could be because some individuals were not exposed to infectious mosquito bites due to spatial heterogeneity or because behavioural factors prevented them from getting bitten, which would not be captured in our model of a homogeneously mixing population. Fitting a model that included a factor to reflect this produced qualitatively the same results as the original model while lowering the reproduction number of dengue in Yap and increasing the proportion estimated to be initially susceptible to dengue infection in Yap as well as the reporting proportion of cases of Zika that were reported. Lastly, the discrepancy could be because some of the population was protected from infection because of cross-immunity from prior infection with another virus, although current evidence points to an opposite effect of antibody-dependent enhancement due to prior dengue infection [48,49]. In the model fits with in this scenario, only the proportion of cases of Zika reported increased. In all cases, this proportion remained well below the equivalent number for dengue.

The case series for Zika in Yap could be interpreted to consist of two peaks. In our basic model, we did not include a mechanism that could have produced these peaks, as we did not have access to any (for example, spatially resolved) data that could have informed such a choice. Whilst two peaks could be produced by a model with spatial heterogeneity, this would have been expected to produce a similar pattern in the dengue outbreak, which consisted of a single peak. Because this is not the case, fits with a two-patch model still yielded a single peak for Zika on the Yap Main Islands. Fitting the Zika outbreak on the Yap Main Islands in isolation using a two-patch model did reproduce two peaks, but ignored the additional information contained in the dengue outbreaks, giving less credence to the fits. In this context, it is worth noting that our model is deterministic and ignores any underlying stochasticity that may have played a role especially early and late in the outbreaks. All uncertainty in our model is in the likelihood which encodes the reporting process. The beginning of what could be seen as a second wave coincided with the arrival of the US Centres for disease Control and Prevention (CDC) teams in Yap, which may have changed reporting rates [9]. With this in mind, our estimate of the proportion of cases reported should be interpreted as an average over the whole outbreak.

Our estimates of human-to-human reproduction numbers for dengue in the Yap Main Islands are consistent with those previously reported in the literature [50], and overlap with the range of 2.8–12.5 estimated from the exponential growth rate alone [51]. The estimate of the human-to-human reproduction number for dengue in Fais, on the other hand, is one of the largest ever observed in the literature, and larger than previous estimates of dengue on small islands [52]. It is conceivable that in this outbreak, everybody was infected within a generation or two. The outbreak hit a population that occupies a small island and is not believed to ever have been exposed to dengue previously, which would explain the rapid spread.

More generally, the estimates for *R*_0_ are similar between dengue and Zika where they have been observed in the same setting on the Yap Main Islands, but differ strongly between the dengue outbreaks on the Yap Main Islands and Fais. This suggests that outbreak setting and human population and mosquito densities are more important in governing transmission dynamics than differences between the pathogens. In other words, while our results suggest that insights from studying dengue transmission in one location can be used to predict the spread of Zika, care must be taken when extrapolating from insights on either of the pathogens in one location to another. Our results suggest that measuring mosquito densities and biting exposure in different settings could provide important information for estimating expected attack rates. In our case, Fais is a much smaller island, and one in which the assumption of random mixing is much more justified than on the Yap Main Islands, where spatial transmission dynamics may dilute the potential for rapid spread, leading to a smaller effective biting rate.

Our estimates of the reproduction number should be interpreted with caution as they could be influenced by heterogeneity. It has been shown if mixing is proportionate but heterogeneous (which is to be expected for dengue or Zika), the reproduction number increases the stronger the heterogeneity [53]. This can cause difficulties in the interpretation of reproduction numbers based on homogeneous models applied to outbreak data [54]. This and other structural limitations of the modelling approach could be contributing in an unknown way to differences or similarities in the estimated values of the reproduction number, and experiments and observational studies will be required to corroborate our findings.

In summary, we have studied three island outbreaks of vector-borne disease and elucidated on similarities and differences. We found that Zika transmission dynamics are similar to dengue when observed in the same setting, and that differences in human population structure and vector density are more important in determining transmission dynamics than difference between the two pathogens. For a new and yet understudied virus such as Zika, comparative studies like this one, especially when conducted on outbreaks in closed populations, can yield important insights into analogies that could be explored for interpreting observed transmission patterns and predicting future dynamics. Field studies on differences in vector density and biting exposure, as well as comparative modelling studies in other settings, would yield important further insights into the relationship between the transmission dynamics of Zika and dengue, and the specific setting in which they occur.

## Acknowledgments

We thank Michael Mina for pointing out an error in an earlier version of the manuscript.

## Supporting Information

### S1 Text

**Modelling results.** R code to reproduce the modelling results, including additional models considered as sensitivity analysis.

### S2 Data

**Time series of cases.** Incidence is given as the number of new cases reported in the week beginning at *onset_week.*

